# Transforming chemical proteomics enrichment into high-throughput method using SP2E workflow

**DOI:** 10.1101/2022.01.24.477214

**Authors:** Tobias Becker, Andreas Wiest, András Telek, Daniel Bejko, Anja Hoffmann-Röder, Pavel Kielkowski

## Abstract

Protein post-translational modifications (PTMs) play a critical role in the regulation of protein catalytic activity, localization and protein-protein interactions. Attachment of PTMs onto proteins significantly diversifies their structure and function resulting in so-called proteoforms. However, the sole identification of post-translationally modified proteins, which are often cell type and disease specific, is still a highly challenging task. Sub-stoichiometric amounts and modification of low abundant proteins necessitate purification or enrichment of the modified proteins. Although the introduction of the mass spectrometry-based chemical proteomic strategies have enabled to screen protein PTMs with increased throughput, sample preparation has remained highly time consuming and tedious. Here, we report an optimized workflow for enrichment of PTM proteins in 96-well plate format which can be possible extended to robotic automatization. This platform allows to significantly lower the input of total protein, which opens up the opportunity to screen specialized and difficult to culture cell lines in high-throughput manner. The presented SP2E protocol is robust, time- and cost-effective as well as suitable for large-scale screening of proteoforms. Application of the SP2E protocol will thus enable the characterization of proteoforms in various processes such as neurodevelopment, neurodegeneration and cancer and may contribute to an overall acceleration of the recently launched Human Proteoform Project.

## Introduction

Protein PTMs are crucial for regulation and fine-tuning of many important biological processes such as neurodevelopment (Becker *et al*, 2021; Mansfield & Gordon-Weeks, 1991; Zheng *et al*, 2021; Aebersold *et al*, 2018), circadian-clocks (Brüning *et al*, 2019) or ageing (Truttmann *et al*, 2018) and impaired in numerous diseases (Rogowski *et al*, 2010; Hoch & Polo, 2020; Kam *et al*, 2018). The incredible diversity of genetic polymorphism, RNA splice variants and PTMs results in many so-called proteoforms (Aebersold *et al*, 2018; Smith *et al*, 2019, 2021), which exceed the ~20,000 human genes by approximately fifty times. This biological network orchestrates the most complex processes including the brain development and ensures a dynamic response of the cells to an external stimulus. However, the extent of protein PTMs in laboratory-cultured cells can differ significantly depending on cell types, diseases and culture conditions. Mass spectrometry (MS)-based chemical proteomics has allowed to reliably map protein PTMs across various experimental conditions (Laughlin & Bertozzi, 2007; Kallemeijn *et al*, 2021; Martin *et al*, 2011; Parker & Pratt, 2020; Kielkowski *et al*, 2020a, 2020b). A widespread application of the chemical proteomic strategy was enabled by parallel improvements of liquid chromatography technologies, gains in speed and sensitivity of mass spectrometers and bioinformatic pipelines for protein identification and quantification (Sinha & Mann, 2020; Cox & Mann, 2008; Tyanova *et al*, 2016; Yu *et al*, 2020). Nowadays, chemical proteomics is used to uncover the scope of protein PTMs in different cell types by the development of small molecule probes which mimic their natural counterparts. The utilization of these probes has provided valuable insights into protein acetylation, palmitoylation, myristylation, prenylation, glycosylation, ADP-ribosylation and AMPylation (Grammel *et al*, 2011; Becker *et al*, 2021; Kliza *et al*, 2021; Laughlin & Bertozzi, 2007; Martin *et al*, 2011; Kallemeijn *et al*, 2021; Yang *et al*, 2010; Sieber *et al*, 2020). In general, different chemical proteomic workflows follow the same sequence of the key steps (Fig 1A). First, cultured cells are treated with the probe which infiltrates the cellular system and competes with the endogenous substrate for the active site of the PTM writer enzymes. Next, the chemical proteomic probes usually contain an alkyne or azide handle to facilitate a bioorthogonal coupling to suitable biotin linkers with either Cu-catalyzed alkyne-azide cycloaddition (CuAAC) or copper free strain-promoted azide-alkyne cycloaddition (SPAAC), respectively (Parker & Pratt, 2020; Agard *et al*, 2004). Following the click chemistry, proteins are precipitated from to remove the excess of biotin reagents and non-protein components of the cell lysate (Kallemeijn *et al*, 2021; Becker *et al*, 2021). In the next step, biotin-labelled proteins are enriched using avidin-coated beads. The critical part of this step is to maximize the efficiency of the washing to remove nonspecifically bound proteins and thus to reduce the complexity of the final MS sample. After reduction and alkylation of the enriched modified proteins, they are digested by trypsin or another protease, desalted and concentrated for MS measurement. The measurement time can be lowered by multiplication using plethora of isotopically labelled MS tags such as tandem mass tag (TMT)(Zecha *et al*, 2019). Alternatively, each sample is measured separately for the label free quantification (LFQ) providing the possibility to add more samples into dataset later on (Cox *et al*, 2014). MS data are typically acquired on orbitrap or timsTOF-based LC-MS/MS instruments. Finally, peptide and protein identifications and quantifications are carried out using well established commercial or free of charge pipelines such as MaxQuant or MSFragger (Yu *et al*, 2020; Cox *et al*, 2014). Comparison of the probe treated and control cells allows to distinguish unspecific protein binders and probe modified proteins. Despite the success of the chemical proteomic technology, the community of scientists combining organic synthesis, mass spectrometry and biology is still rather small. With numerous validated commercial PTM probes and the widespread availability of mass spectrometers either used in individual groups or as a core services, the bottlenecks of the chemical proteomic approach still remain in the insufficient consistency, time inefficiency and laboriousness of the enrichment techniques. Furthermore, the emerging field of chemical proteomic studies focused on neuronal differentiation and the complex environment of the central nervous system composed of many different cell types, is not compatible with the high amounts of total proteins that have been so far required for the analysis. In a perfect scenario, the workflow would be efficient even with a low protein input protein and the enrichment would require a minimum hands-on time or fully automatization. In comparison to standard MS whole proteome sample preparation methods, which include filter-assisted sample preparation (FASP), stage tips, iST, and single-pot solid-phase amplified sample preparation (SP3), the chemical proteomic method not only requires highly efficient protein and peptide purification, but also needs to be combined with suitable bioorthogonal reaction conditions and much higher starting protein amounts (Wiśniewski *et al*, 2009; Müller *et al*, 2020; Hughes *et al*, 2019; Sielaff *et al*, 2017). Thus far, the most common chemical proteomic methods for protein PTM enrichment utilize acetone or chloroform-methanol precipitation to remove the excess of click chemistry reagents (Kallemeijn *et al*, 2021). Further, enrichment with avidin-coated agarose beads requires centrifugation or filtration to separate them from the wash buffer. Of note, the Tate group combined avidin-coated magnetic beads and a trifunctional linker with azide, biotin and rhodamine to visualize the enriched proteins by in-gel analysis (Wright *et al*, 2014; Kallemeijn *et al*, 2021) and recently the Backus group has implemented the SP3 peptide cleanup into their chemical proteomic workflow before transferring the peptides including the biotin-modified peptides on avidin-coated agarose beads (Yan *et al*, 2021). Although similar affinity enrichment techniques combined with MS have been reported, to our best knowledge the procedure feasibly integrating all aspects of small-scale chemical proteomics is not available (Klont *et al*, 2021; Makowski *et al*, 2018).

**Figure 1.**
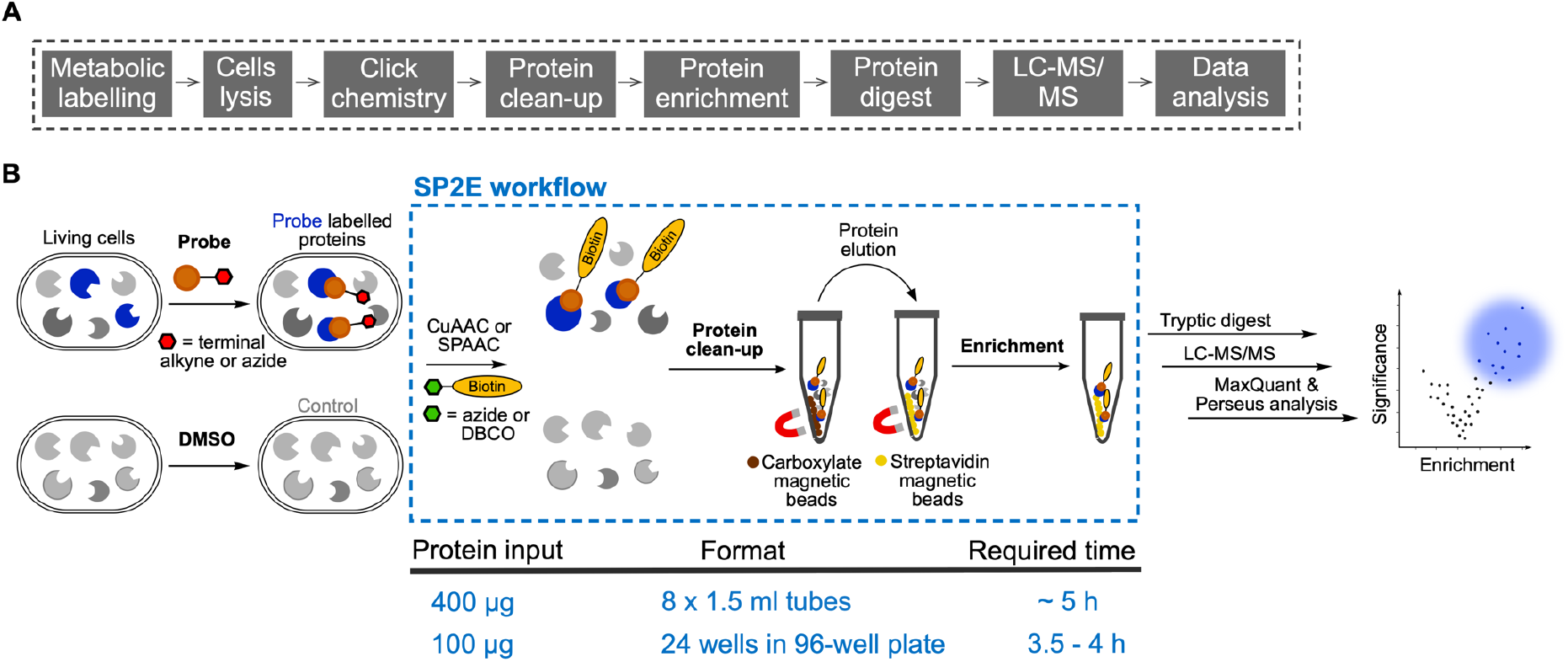
Schematic overview chemical proteomic workflow. A Standard chemical proteomic workflow key steps. B Schematic characterization of the SP2E workflow and basic parameters of the procedure.

Here, we report the development and optimization of the chemical proteomic method which uses carboxylate-modified magnetic beads to clean-up the proteins after CuAAC and streptavidin-coated magnetic beads for the enrichment of the labelled proteins (Fig 1B). The new method termed SP2E was further scaled down to 96-well plate format, starting with 100 μg total protein. The SP2E method has been successfully used for profiling of protein glycosylation and the low abundant protein PTM called AMPylation. Together, the SP2E method provides a time-effective and robust platform for routine and high-throughput profiling of protein PTMs.

## Results

### Development of the SP2E workflow for chemical proteomics

First, we set to optimize the lysis buffer composition to maximize the efficiency of the click chemistry. To evaluate this, the HeLa cells were treated with the pro-N6pA probe infiltrating protein AMPylation (Becker *et al*, 2021; Kielkowski *et al*, 2020b), harvested and lysed in nine different lysis buffers (Fig 2A). The CuAAC click chemistry was performed with 400 μg total protein per sample and azide-TAMRA to visualize the conversion efficiency by in-gel fluorescence scanning after sodium dodecyl-sulfate polyacrylamide gel (SDS-PAGE) electrophoresis. The overall brightest fluorescence was observed in the lysis buffer containing 0.1% NP-40, 0.2% SDS in 20 mM Hepes pH 7.5 (Fig 2B), which was used for all following experiments. Next, we focused on optimization of protein enrichment and mass sample preparation. To assess the enrichment efficiency, we decided to use a group of six known AMPylated maker proteins (HSPA5, CTSB, PFKP, PPME1, ACP2, ABHD6). In the first attempt 400 μg total protein in 200 μL lysis buffer was used for click chemistry with the azide-PEG_3_-biotin. The resulting reaction mixture was transferred onto carboxylate-coated magnetic beads and followed by the addition of absolute EtOH to a final concentration of 60%. After washing the beads three times with 80% EtOH, the streptavidin-coated magnetic beads were added directly to the carboxylate-coated magnetic beads and incubated for 1 h in 0.2% SDS in PBS to form the biotin-streptavidin complex. To remove the unmodified proteins, the beads mixture was washed thrice with 0.1% NP-40 in PBS, twice with 6M urea in water and thrice with LC-MS grade water. The enriched proteins were subsequently reduced, alkylated and trypsin digested in ammonium bicarbonate (ABC) buffer. The resulting peptides were eluted from the beads by two washes with the ABC buffer before desalting on off-line Sep-Pak C18 columns and separation on a UHPLC using a 150 min gradient with the high fidelity asymmetric waveform ion mobility spectrometry (FAIMS) device attached on the Orbitrap Eclipse Tribrid mass spectrometer. The MS data were analyzed by MaxQuant and evaluated in Perseus (Fig 2C)(Cox *et al*, 2014; Tyanova *et al*, 2016).

**Figure 2.**
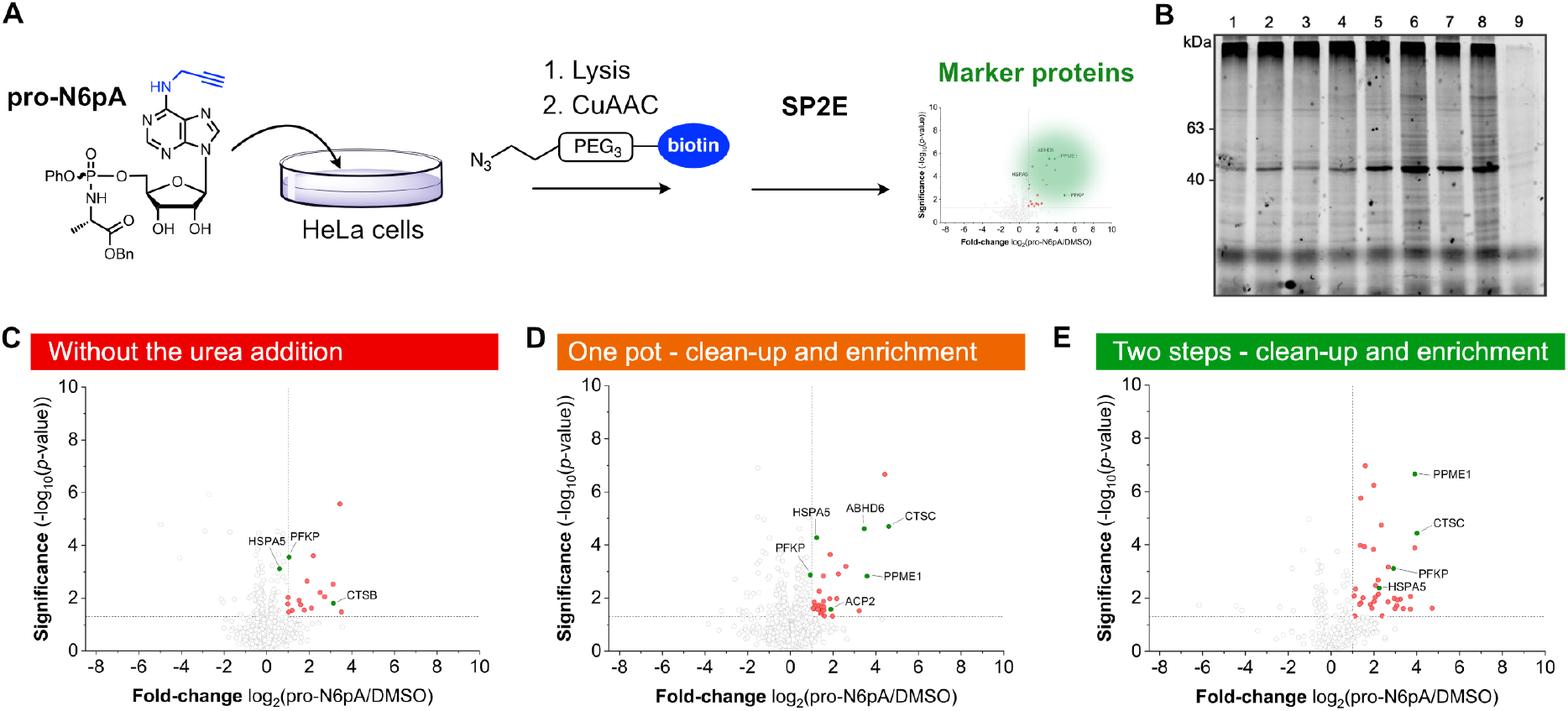
Development and optimization of the SP2E workflow using AMPylation probe. A Pro-N6pA probe structure and workflow used for the optimisation of the SP2E method. B Optimization of the lysis buffer based on the efficiency of the CuAAC click chemistry. Lysis buffer compositions: line 1 (1% NP-40 in PBS), line 2 (1% NP-40, 0.2% SDS in PBS), line 3 (0.5% Triton in PBS), line 4 (0.5% Triton, 0.2% SDS in PBS), line 5 (1% NP-40 in 20mM Hepes), line 6 (1% NP-40, 0.2% SDS in 20mM Hepes), line 7 (0.5% Triton in 20mM Hepes), line 8 (0.5% Triton, 0.2% SDS in 20mM Hepes), line 9 (8M urea in 0.1M Tris/HCl). C Volcano plot showing significantly enriched proteins (red dots) using the pro-N6pA AMPylation probe with highlighted marker proteins (green dots). The proteins were loaded onto carboxylate magnetic beads in the lysis buffer and enrichment was done in one pot combining carboxylate and streptavidin magnetic beads together. D The proteins after click reaction were loaded in lysis buffer supplemented with 4M urea. The enrichment was done in one pot together with carboxylate magnetic beads. E Compared to the protocol used for Fig. 2C, the proteins were eluted after the protein cleanup on carboxylate coated magnetic beads and were transferred into a new 1.5 mL tube for enrichment containing streptavidin magnetic beads. All volcano plots; *n* = 4, cut-off lines at *p*-value >0.05 and 2-fold enrichment.

We have observed that protein loading in lysis buffer directly after the click reaction onto carboxylate magnetic beads did not give satisfactory results with poor enrichment of the marker proteins (Fig 2C). Therefore, based on our previous experiments, we have tested whether the addition of concentrated urea to the click reaction mixture in lysis buffer may improve the protein clean-up. Indeed, the dilution of the lysis buffer to a final concentration of 4M urea has significantly improved the overall enrichment ratio (Fig 2D).

In the second step, we asked, whether it is possible to reduce the overall background by eluting the proteins from the carboxylate magnetic beads before adding them to the streptavidin-coated magnetic beads in a new tube, and thus improve the enrichment ratio. Therefore, directly after the clean-up on carboxylate magnetic beads, the proteins were eluted twice with 0.2% SDS in PBS and transferred onto streptavidin magnetic beads. The resulting volcano plot has confirmed that separate elution and transfer of the proteins are beneficial for their enrichment and hence outcompetes the advantage of performing both steps in one-pot (Fig 2E). In particular, the enrichment of HSPA5 and PFKP has increased by more than 2-fold.

Third, the digest and peptide elution conditions have been optimized. In principle, the possibility to combine the click chemistry with reduction and alkylation prior to protein cleanup on carboxylate beads, as described by Hughes *et al*. (Hughes *et al*, 2019, 2014), could improve the final elution and purity of the peptides. However, experimental testing of this possibility showed very poor enrichment of modified proteins, while standard reduction and alkylation with tris(2-carboxyethyl)phosphine (TCEP) and chloroacetamide (CAA) at 95 °C for 5 min directly before trypsin addition gave satisfying results (Fig 2E).

Taken together, we have optimized several steps of the SP2E protocol. First, we showed that the lysis buffer containing 0.1% NP-40, 0.2% SDS in 20 mM Hepes pH 7.5 efficiently lyse the cells and improve the yield of the CuAAC. Furthermore, the addition of urea into the lysis buffer after click chemistry enhances protein binding to the carboxylate magnetic beads. Next, we demonstrated that it is possible to reduce the background by eluting the proteins from the carboxylated magnetic beads before adding them to the streptavidin magnetic beads. Finally, it was confirmed that the reduction and alkylation after the proteins’ enrichment and prior to trypsination worked better in comparison to reduction and alkylation performed before the protein clean-up.

### Application of the SP2E workflow for analysis of protein AMPylation

To validate our approach a heterogeneous set of samples, we went on to screen metabolic pathways which may impact on protein AMPylation. In our previous studies, we showed that changes in protein AMPylation are linked to neurodevelopment of the human induced pluripotent stem cells (hiPSCs) (Becker *et al*, 2021; Kielkowski *et al*, 2020b). In particular, we have observed that a large group of lysosomal proteins is modified with the pro-N6pA probe and thus likely AMPylated. Most recently, we have described the intriguing changes in proteoforms of the 5’-3’ exonuclease PLD3 (PLD3), which leads to accumulation of the soluble modified PLD3 in mature neurons. In line with our marker proteins used for the SP2E workflow development, we have consistently enriched a group of proteins localized to cytosol and mitochondria such as PFKP, PPME1, SLC25A3 and cytoskeletal proteins. Therefore, our and other reports has led to a controversy in the field, as the two thus far known AMPylators, FICD and SELENOO, are strictly localized to the endoplasmic reticulum (ER) and mitochondria, respectively (Truttmann *et al*, 2018, 2017; Sanyal *et al*, 2019). Now, with our optimized SP2E protocol in hand, we decided to determine the signaling pathways in which AMPylation may play a role. We have proceeded with a screening of five different inhibitors in SH-SY5Y neuroblastoma cells: rapamycin, bafilomycin, 2-deoxy-D-glucose, thenoyltrifluoroacetone (TTFA) and monensin. These inhibitors regulate mTOR, autophagy, glycolysis, cellular respiration and ER to Golgi sorting, respectively (Raught *et al*, 2001; Leeman *et al*, 2018; Yoshimori *et al*, 1991; Zhang & Fariss, 2002; Mollenhauer *et al*, 1990). SH-SY5Y cells were treated either with the inhibitor alone or with the inhibitor and pro-N6pA probe to avoid any influence of protein expression changes triggered by addition of the inhibitor (Fig 3A). Additionally, two more controls were included. Cells treated with plain DMSO or only pro-N6pA probe to ensure the consistency with the previous results and to check the efficiency of the enrichment (Fig 3B). For each condition, four replicates have been prepared and the enrichment protocol has been started with 400 μg total protein (Fig 3A). The resulting MS samples were analysed using a LC-MS/MS 150 min gradient and the peptides were identified and quantified by the LFQ method in MaxQuant. This has resulted in overall 100 significantly enriched proteins. The principal component analysis (PCA) showed a clear difference between control and probe treated samples and, interestingly, the samples that were treated with bafilomycin and monensin clustered together suggesting the robustness and feasibility of the SP2E workflow to discover new pathways involved in regulation of AMPylation (Fig 3C and Fig S1 and S2). Pearson correlation coefficients of MS intensities between the replicates has been over 95% (Fig 3D). Of note, the analysis of the enriched proteins revealed the amyloid-beta precursor protein (APP) to be one of the most significantly enriched protein in cells treated with bafilomycin and monensin.

**Figure 3.**
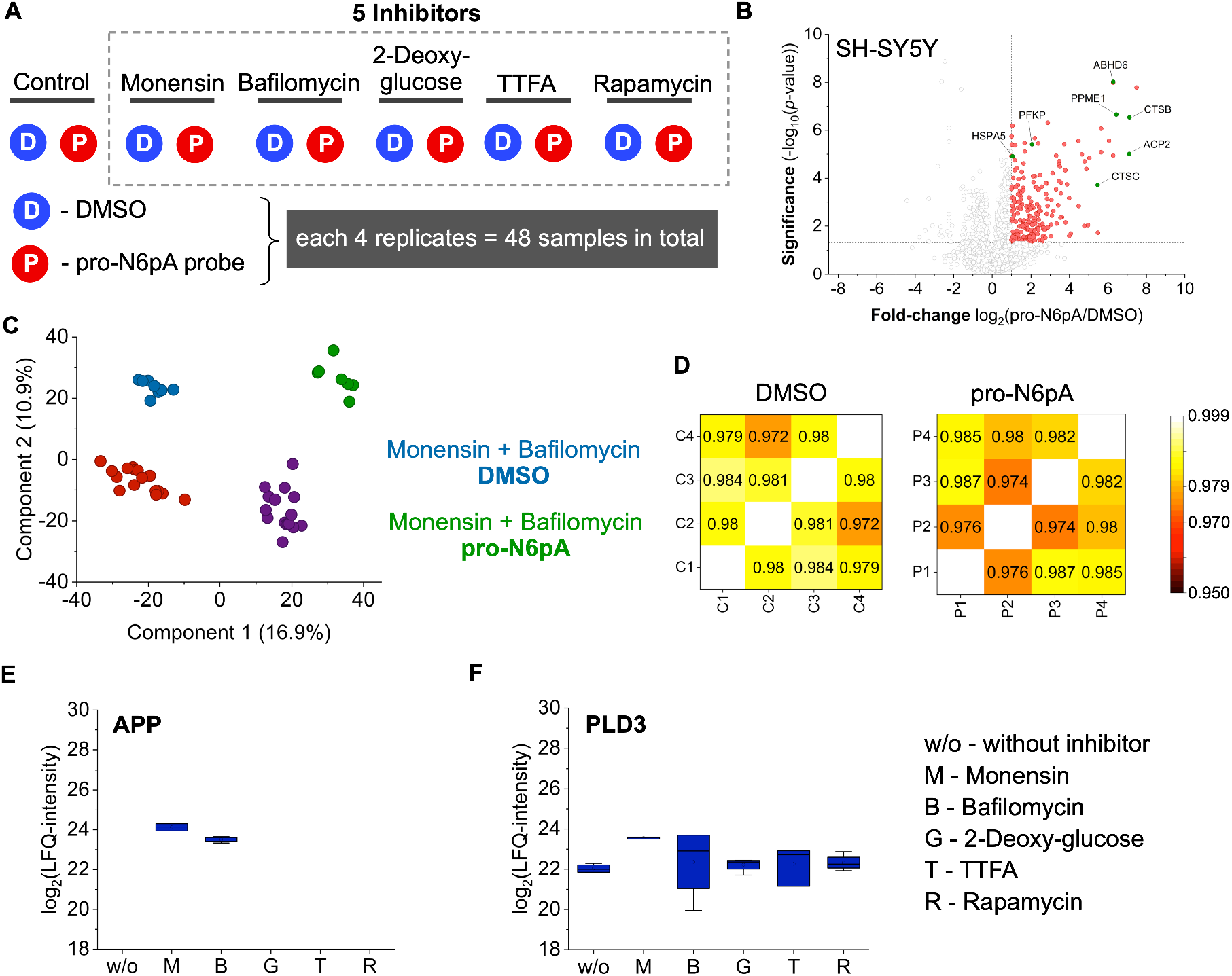
Analysis of protein AMPylation under different stress conditions using the SP2E workflow. A Design of the experiment to test the impact of various inhibitors on protein AMPylation. B Volcano plot showing the enrichment of AMPylated proteins (pro-N6pA vs. DMSO) from SH-SY5Y cells using the SP2E protocol; *n* = 4, cut-off lines at *p*-value >0.05 and 2-fold enrichment. C PCA of the inhibitors treated cells and controls display separation of the monensin and bafilomycin as well as pro-N6pA treated cells. D Representative heatmaps visualizing the Pearson correlation coefficients of LFQ intensities of DMSO and pro-N6pA replicates. E Profile plot displays the APP LFQ intensities under various conditions. The APP was not found in any other conditions, for example in cells only treated with DMSO or inhibitor. F Profile plot displays the PLD3 LFQ intensities under various conditions.

Next, examination of the profile plots without imputed values showed that APP is only enriched in the cells that were treated with the pro-N6pA probe and either bafilomycin or monensin (Fig 3E). Interestingly, PLD3 has shown a strong response to the two inhibitors (Fig 3F). Subsequent search for similar enrichment profiles has uncovered a group of 12 proteins including GPR56, FAT1, LAMA4, TGOLN2, RNF149, CRIM1, ITM2B, L1CAM, TMEM59, MCAM, LRP1 and CLU that were specifically enriched under these two conditions. This would point towards the link between AMPylation and trafficking pathways from ER to lysosomes and autophagy.

To investigate the relationship between the monensin concentration and the AMPylation extent in more detail, SH-SY5Y cells were treated with increasing concentration of monensin in cell culture media ranging from 2 nM to 2 μM (Fig 4A). The concentration of the pro-N6pA probe was kept constant and each condition has been performed in duplicate. Surprisingly, the modification of PLD3 has risen by several folds even with the lowest inhibitor concentration and then it was steadily increasing with the monensin concentration (Fig 4B). In contrast, the modified APP was only found in cells treated with 1 μM and 2 μM monensin (Fig 4C). Indeed, PTMs of the amyloid-beta precursor protein are deemed to play an important role in development of the Alzheimer’s diseases pathophysiology(Long & Holtzman, 2019). By application of our SP2E workflow we showed that AMPylation might be an additional PTM involved in the regulation of the APP physiological function. Together, the screening of the AMPylation changes triggered by five different active compounds manifest the utility of the SP2E workflow, which is characterised by minimal background binding, its robustness and replicability.

**Figure 4.**
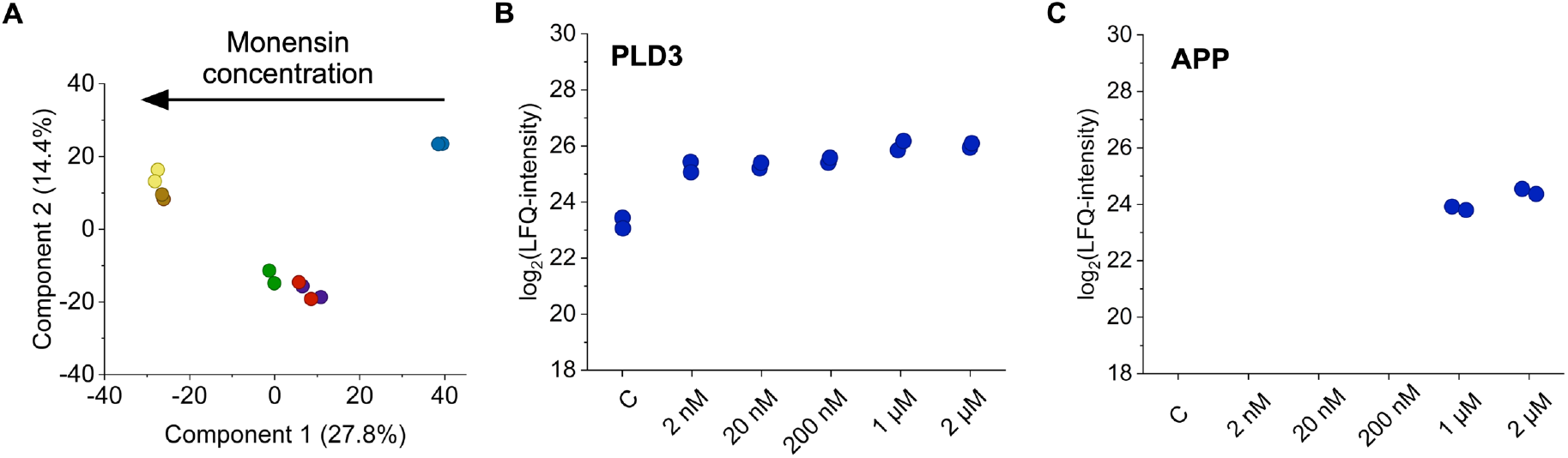
Monensin concentration dependent increase in APP and PLD3 modification. A PCA displays distinct changes in the enriched proteins with increasing monensin concentration. B Profile plot of the PLD3 LFQ intensities shows rapid increase of the PLD3 modification with 2nM monensin concentration. C In contrast to PLD3, the profile plot of the APP LFQ intensity reveals that APP is only enriched with the pro-N6pA probe with 1 μM and 2 μM monensin in cell culture media.

### Application of the SP2E workflow for analysis of protein *O*-GlcNAcylation

To assess the utility of our optimized workflow for other PTMs, we used the previously described Ac_3_4dGlcNAz probe (GlcNAz) for the pull-down of *O*-linked β-*N*-acetyl-glucosamine glycosylated (*O*-GlcNAcylation) proteins (Fig 5A)(Yang & Qian, 2017; Laughlin & Bertozzi, 2007; Li *et al*, 2016; Pedowitz & Pratt, 2021). Numerous metabolic labels have been developed for characterisation of *O*-GlcNAcylated proteins (Li *et al*, 2016). However, they often suffer from low substrate specificity and labelling efficiency. Here, we used the 2,4-dideoxy-D-glucopyranose derivative, which shows improved specificity for cytosolic proteins (Li *et al*, 2016). In contrast to previous experiments with the pro-N6pA probe, the GlcNAz probe contains an azide functional group for bioorthogonal protein labelling using SPAAC (Fig 5A)(Agard *et al*, 2004). To avoid unspecific reactivity of free thiols with the DBCO-biotin reagent utilized for the SPAAC, they were capped with iodoacetamide (Pentelute *et al*, 2018). Afterwards, the lysate proteins containing GlcNAz were reacted with the DBCO-biotin reagent and enriched using the SP2E method in the same fashion as described above for AMPylation (Fig 5B). Interestingly, the high number of significantly enriched proteins leads to a clear separation of the probe treated and control samples in the PCA plot, with one component over 72 % (Fig 5C). Furthermore, the SP2E enrichment of the *O*-GlcNAcylated proteins had a strong impact on the number of imputed values, as a high number of proteins were consistently identified only in the probe treated samples (Fig 5D). To our contentment, 95% of the 358 significantly enriched proteins were previously described as *O*-GlcNAcylated (www.oglcnac.mcw.edu), with one of the most significant hits being the well-studied co-translationally *O*-GlcNAcylated protein NUP62 (Fig 5B-E, Fig S3)(Wulff-Fuentes *et al*, 2021). Together, these experiments demonstrate the utility of the SP2E protocol for the enrichment of metabolically labelled proteins.

**Figure 5.**
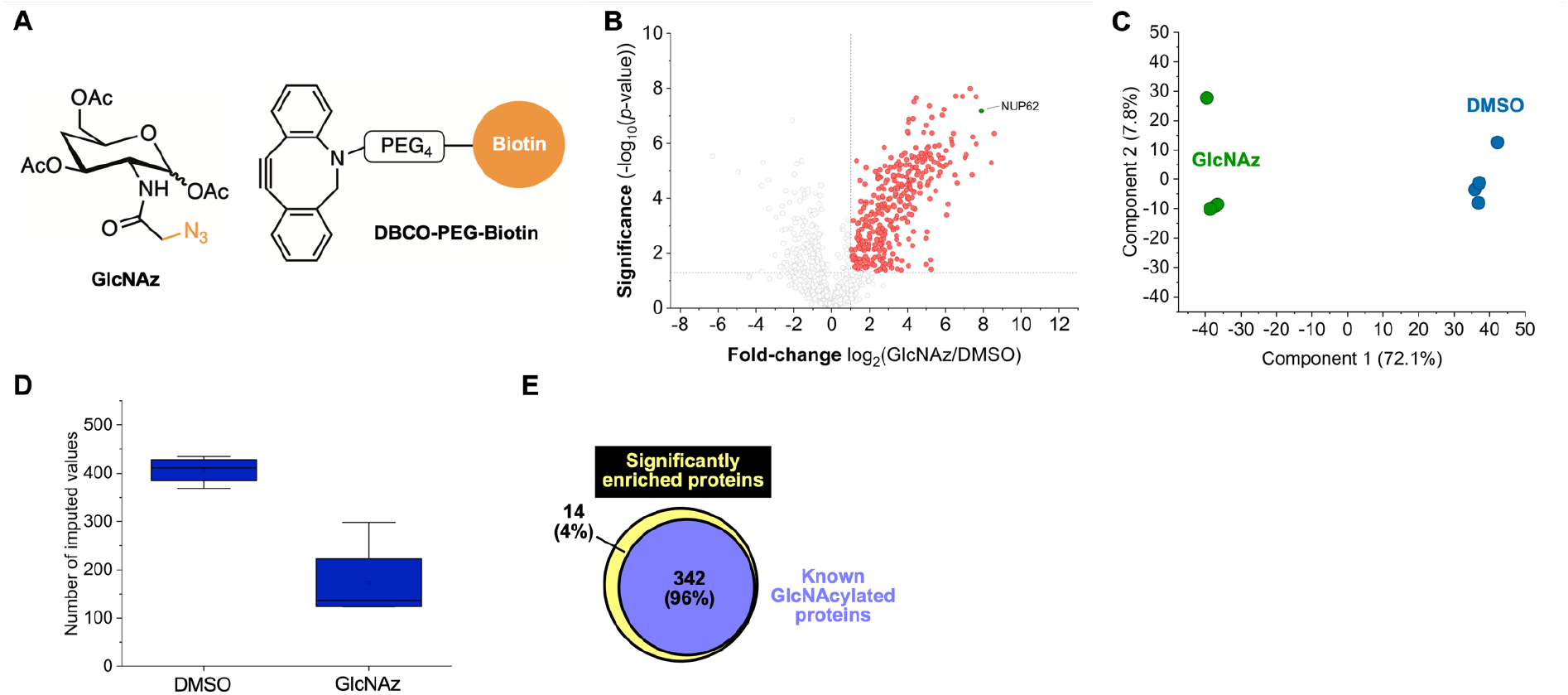
Analysis of O-GlcNAcylation by GlcNAz probe, SPAAC and SP2E workflow. A Chemical structure of GlcNAz probe for metabolic labelling of *O*-GlcNAcylated proteins and DBCO-biotin reagent to functionalize the probe modified proteins by SPAAC. B Volcano plot visualizing the enrichment of the *O*-GlcNAcylted proteins; *n* = 4, cut-off lines at *p*-value >0.05 and 2-fold enrichment. Red dots are significantly enriched proteins. C PCA graph points to a clear separation of the control and probe treated samples. Of note, the component 2 possess a high value of 72.1 %. D Comparison of the number of imputed values in DMSO and GlcNAz treated cells to demonstrate the clear difference between the two samples and enrichment efficiency. E Diagram showing overlap between all significantly enriched proteins using GlcNAz probe and previously described *O*-GlcNAz proteins.

### Scale-down of the SP2E workflow into 96-well plate format

Although, the combination of the carboxylate and streptavidin magnetic beads using SP2E streamlined the PTM protein enrichment, it remained to be demonstrated whether this approach is efficient with a lower protein inputs. Therefore, we have performed the enrichment starting with 100 μg of total protein using lysates from pro-N6pA treated HeLa cells in a standard 1.5 mL eppendorf tube. Already in this first attempt, it was possible to significantly enrich 5 out of 6 AMPylation marker proteins (Fig 6A). Encouraged by the general feasibility of the SP2E workflow with lower protein input, we have moved on to scale down the protocol into 96-well plate format. The dynamic range of the SP2E enrichment efficiency has been shown on the enrichment of rather low abundant AMPylated proteins from pro-N6pA treated HeLa cells in 96-well plate format (Fig 6B). The initial testing with simple decrease of the click reaction volume to 20 μL and wash steps to 150 μL showed only poor enrichment results (Fig S4). Thus, we have adjusted the protocol with the following steps. The clean-up of the proteins after the click reaction was extended with an additional acetonitrile washing step, as used for the automated whole proteome samples preparation by Müller et al. (Müller *et al*, 2020). In addition, the reduction and alkylation step were omitted, proteins were digested in 50 μL of TEAB and peptides were eluted from the streptavidin magnetic beads with 20 μL TEAB and 20 μL 0.5% formic acid (FA) buffer with an incubation at 40 °C for 5 min (Fig S4). The resulting MS samples have been acidified by addition of FA and the peptide mixture was resolved using a 60 min LC-MS/MS measurement (Fig 6B). In particular, usage of the shorter gradient is beneficial for two practical reasons. First, more samples can be measured in shorter time and second, the MS spectra files are smaller with overall less MS data to process, leading to a faster identification and quantification by search engines. Furthermore, the total amounts of enriched peptides are estimated to be still very low, the shorter gradient is likely resulting in more intense MS spectra and thus in more identified peptides (Fig 6B). Importantly, the Pearson correlation coefficients of the protein intensities remained still over 95 % (Fig 6C).

**Figure 6.**
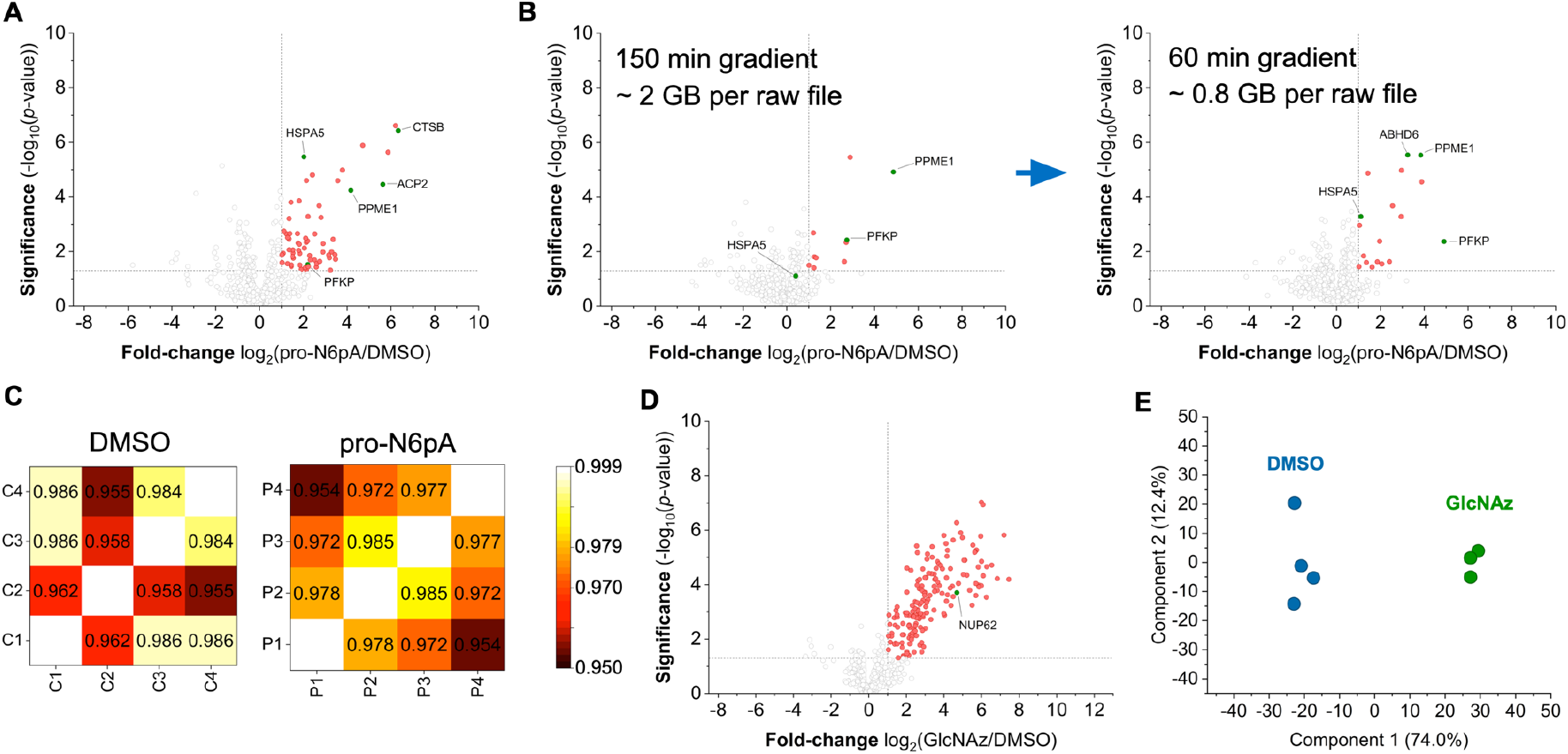
Scale-down of the SP2E workflow into 96-well plate format. A SP2E protocol with 100 μg input protein performed in 1.5 mL tubes visualized in the volcano plot. B Optimization of the LC-MS/MS measurement with 100 μg protein input using the 96-well plate format SP2E protocol. C Heatmaps representing the Pearson correlation coefficients between the replicates. D Volcano plot showing the enrichment of *O*-GlcNAcylated proteins starting from 100 μg input protein in 96-well plate format. E PCA of the small scale GlcNAz enrichment shows a slightly better separation of controls from probe treated samples compared to large-scale SP2E in Fig 5B. All volcano plots, *n* = 4, cut-off lines at *p*-value >0.05 and 2-fold enrichment. Red dots are significantly enriched proteins.

The efficiency of the optimised protocol in a 96-well plate has been further tested with the GlcNAz probe (Fig 6D). Indeed, it was possible to significantly enrich 174 proteins with NUP62 amongst the most significantly enriched ones (Fig 6D). Moreover, both TEAB and ABC buffers used for the digest provided comparable results. In order to elute the peptides efficiently from the streptavidin magnetic beads after digest, it is necessary to repeat the elution twice, although it results in peptides dilution in the final MS sample (Fig S4). Similar to the large-scale experiment, the SP2E procedure in the 96-well plate format displays an excellent separation in the PCA of the control and probe treated samples after enrichment (Fig 6E). In summary, our protocol promises to provide a fast, robust and high-throughput chemical proteomic platform, which may be used by biologists to assess the PTM status from a wide variety of cells. This is an important prerequisite to unravel the complex PTM networks and elucidating the underlying functional consequences of protein PTMs.

## Discussion

Chemical proteomics has enabled the characterization of many protein PTMs, which are otherwise inaccessible using the whole proteome analysis. Several enrichment workflows have been developed to make the procedure universal and feasible. However, the protocols often require to be carried out by specialized laboratory personnel, they are tedious and time consuming. Additionally, with increasing demand to screen protein PTMs in specialized cell types which are difficult to culture or not accessible in larger amounts typically required by the chemical proteomic protocols, it is of the paramount importance to streamline the enrichment workflow and to provide a platform which could be automatized. This would parallel the development of the high-throughput automatized whole proteome MS analysis. The transformation of the MS field has been underlined by the rapid improvement of speed and sensitivity of nowadays mass spectrometers. The progress of the MS instrumentation has been complemented by software tools allowing for fast and reliable protein identification and quantification. Together, these developments have created the suitable environment for transition of the chemical proteomic analysis of protein PTMs from the specialized field to become a more widely applied analytical tool.

Here, we describe the development and application of the SP2E workflow which enables the chemical proteomic characterization of protein PTMs in a small-scale, robust and time effective manner. The main difference to the previous MS-based chemical proteomic protocols is the utilization of the paramagnetic beads for both protein clean-up and enrichment. It leads to a better separation of solid and liquid phase and thus improved the removal of nonspecifically binding proteins during the enrichment steps. Furthermore, it allows a better separation as well in smaller volumes and it can be readily automatized. The initial substitution of the standardly used avidin-coated agarose beads with streptavidin-coated magnetic beads resulted in only moderate enrichment of the AMPylated marker proteins. Therefore, we have systematically evaluated each step of the enrichment protocol with focus on scale-down of the whole procedure. Starting with the lysis buffer, which is critical to ensure an efficient lysis already in a small volume to provide protein concentrations of up to 10 μg/μL and efficient click chemistry (Fig 2A). To complete the optimization of the Cu-catalyzed click reaction time, a time dependent experiment was carried out to show that the 1.5 h is necessary to maximize the yield of the reaction (Fig 7A). Next, enrichment efficiency was improved by separation of protein clean-up and enrichment which possibly allow efficient washing and, hence, less unspecific protein binding. The optimized protocol has been tested in large-scale experiments starting with 400 total protein to explore metabolic pathways in which protein AMPylation plays a role. In total, 48 samples have been prepared using five different inhibitors and four replicates per condition. The same pro-N6pA probe was utilized as for the optimisation of the workflow with the exception of SH-SY5Y neuroblastoma cells instead of the HeLa cells were used. Pearson correlation coefficient of protein intensities among all samples showed high correlation (> 0.95), demonstrating the robustness of the workflow. Interestingly, the PCA revealed a difference between the control and probe treated cells and importantly displays a clear change in enriched proteins from monensin and bafilomycin treated cells suggesting the specific role of AMPylation in cell stress response to these inhibitors. Moreover, this indicates the high efficiency of the wash steps during enrichment and reproducibility of the SP2E workflow. Concentration dependent analysis of monensin on pro-N6pA labelling showed that already two replicates of each condition provide sufficient information due to the high reproducibility of the procedure and revealed that the PLD3 modification rapidly increases with 2 nM monensin concentration. This was then confirmed in the follow up in-gel analysis of the PLD3 using a trifunctional linker containing an azide, biotin and rhodamine. After SP2E based enrichment, proteins are released from the streptavidin magnetic beads by denaturation and separation on SDS-PAGE. Then, enrichment efficiency is assessed by in-gel fluorescence. Enrichment of PLD3 was additionally confirmed by western blot and, for the first time, uncover for the first time that only the soluble form of PLD3 is modified by the pro-N6pA probe and that there is a strong increase in the modified form upon addition of monensin (Fig 7B). Even though, the soluble and soluble-modified PLD3 are not even visible on the western blot of the whole lysate and the increase in modified PLD3 after enrichment is obvious (Fig 7C, D). Together, this demonstrates the efficiency of the SP2E enrichment.

**Figure 7.**
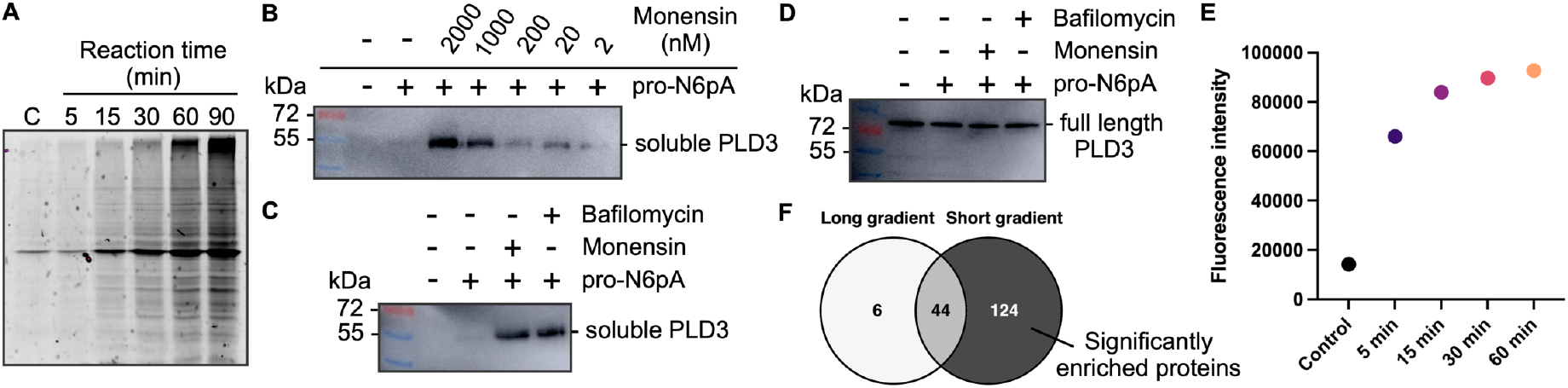
SP2E time-efficiency optimization and in-gel analysis of the PLD3 modification. A In-gel fluorescence showing the click reaction time optimization. In the control (C), cells were treated with plain DMSO and the lysate was incubated with the click reaction mixture for 90 min. B Enrichment of the modified PLD3 using the trifunctional linker (azide-biotin-rhodamine) in dependency on monensin concentration in cell culture media. PLD3 detected on the PVDF membrane using the anti-PLD3 antibody. C Enrichment of the modified PLD3 using the trifunctional linker (azide-biotin-rhodamine) after the treatment with bafilomycin (100 nM) and monensin (2 μM). D Western blot of the whole proteome from cells treated with bafilomycin and monensin stained with anti-PLD3 antibody. E Plot display total fluorescence intensity from the in-gel analysis of time optimization of biotin-streptavidin complex formation (see original gel in Fig S5). F Venn diagram comparing the number of significantly enriched proteins found after 96-well plate format SP2E of GlcNAcylated proteins using short (60 min) or long (150 min) LC-gradient.

To show the versatility of the procedure, we have performed the enrichment of *O*-GlcNAcylated proteins using the azido GlcNAz probe. The SPAAC click reaction followed by the SP2E workflow provided excellent enrichment of the well described glycosylated protein NUP62 with nominal values of 240-fold enrichment and a *p*-value of 1E-7. Furthermore, another 342 known glycosylated proteins were significantly enriched (95.5 %). The outstanding enrichment efficiency is visible by the number of missing values in the control samples (Fig 5D). Since protein glycosylation plays an important role in numerous metabolic processes and often serves as disease marker, our SP2E protocol offers the possibility to screen for the *O*-GlcNAcylation in a high-throughput manner. This might not only accelerate the progress in the field, but also help to decipher the complex glycan patterns by application of different glycosylation labels.

Finally, current availability of chemical proteomic data shared in public repositories together with feasibility of sophisticated data analyses necessitate generation of high-quality data in a high-throughput manner. This can be only achieved by automatization of the procedures as for example with the autoSP3 protocol and other proteomic approaches. Here, we have scaled-down the SP2E workflow into the 96-well plate format, which retains the principle operations paralleled in autoSP3. Importantly, we showed on analysis of protein AMPylation and *O*-GlcNAcylation that 100 μg input protein is sufficient for successful PTM protein profiling. This results in the same high correlations between the samples (in average >0.96) and achieves high fold-enrichments (NUP62 26-fold and HSPA5 2-fold). Although there is a drop in the overall number of enriched proteins, to 48% for AMPylation and 47% for *O*-GlcNAcylation, this is outweighed by cost and time efficiency due to smaller scale of cell culture, washing steps, usage of the multichannel pipettes, shorter measurement times and data processing. Of note, in case of AMPylation, 66% marker proteins were successfully enriched and for *O*-GlcNAcylation 173 proteins out of 174 significantly enriched proteins were previously described *O*-GlcNAcylated. To further shorten the time of the workflow, we have analysed the time-dependency of the biotin-streptavidin complex formation to find out that already 15 min is sufficient (Fig 7E, F respectively). Overall, the manual SP2E workflow with 24 samples in 96-well format can be carried out in 3.5 - 4 hours starting from the preparation of the click reaction to the addition of trypsin. After overnight trypsin digest, the peptides are eluted and transferred into MS-vials within additional 45 min.

The limitation of the SP2E workflow for chemical proteomic characterisation of protein PTMs is the inherent necessity to treat the cells with small molecule probes, which are not always commercially available and thus need to be synthesized. Furthermore, the labelling ratio is often determined by the metabolic activation of probes and the substrate selectivity of PTM writers. Thus, it might be difficult to estimate the exact stoichiometry of protein modification. Similar to other proteomic approaches, the statistical evaluation includes the imputation of missing values and therefore it might be challenging to identify the hit proteins with low protein intensities as significantly enriched. However, some of these bottlenecks can be overcome by increasing the number of replicates.

In summary, the robust, time and cost-effective workflow we have developed will allow a wide range of scientist to characterise protein PTMs in the future, which might accelerate the deconvolution of the function of low abundant protein PTMs in diseases.

## Materials and Methods

### Reagents and Tools Table

| Reagent/Resource | Reference or Source | Identifier or Catalog Number |
| --- | --- | --- |
| **Experimental Models** |  |  |
| Human: SH-SY5Y | Biedler et al., 1973 | RRID: CVCL_0019 |
| Human: HeLa | Puck and Marcus, 1955 | RRID: CVCL_0030 |
| **Recombinant DNA** |  |  |
| / |  |  |
| **Antibodies** |  |  |
| Rabbit polyclonal anti-PLD3 | Sigma-Aldrich | Cat# HPA012800; RRID: AB_1855330 |
| Goat polyclonal anti-rabbit IgG, HRP-linked | Sigma-Aldrich | Cat# A6667; RRID: AB_258307 |
| **Oligonucleotides and other sequence-based reagents** |  |  |
| / |  |  |
| **Chemicals, Enzymes and other reagents** |  |  |
| Acetonitrile (LC-MS grade) | Thermo Fisher Scientific | Cat# A955-212 |
| **Alanyl-Glutamine** | Sigma-Aldrich | Cat# G8541 |
| Ammoniumperoxodisulfat (APS) | Sigma-Aldrich | Cat# 09913 |
| Biotin-PEG<sub>3</sub>-N<sub>3</sub> | Carbosynth | Cat# FA34890 |
| BSA | AppliChem | Cat# A6588 |
| Coomassie Blue R-250 | Fluka | Cat# 27816 |
| CuSO<sub>4</sub> x 5 H<sub>2</sub>O | Acros | Cat# 10627162 |
| ddH<sub>2</sub>O (LC-MS grade) | Honeywell | Cat# 15665350 |
| DMSO | Sigma-Aldrich | Cat# D4540 |
| Formic Acid (LC-MS grade) | Thermo Fisher Scientific | Cat# A117 |
| Hepes | Carl Roth | Cat# HN77.5 |
| NP40 | Sigma-Aldrich | Cat# 74385 |
| Methanol (LC-MS grade) | Thermo Fisher Scientific | Cat# A456 |
| Powdered milk | AppliChem | Cat# A0830 |
| SDS | AppliChem | Cat# A2572 |
| TAMRA-N<sub>3</sub> | Baseclick | Cat# BCFA-008-1 |
| TBTA | TCI | Cat# T2993 |
| TCEP | Carbosynth | Cat# FT01756 |
| TEAB (1 M) | Sigma-Aldrich | Cat# T7408 |
| TEMED | Sigma-Aldrich | Cat# T9281 |
| Tris-base | Thermo Fisher Scientific | Cat# 10724344 |
| Trypan Blue | Thermo Fisher Scientific | Cat# 11538886 |
| Urea | AppliChem | Cat# A1049 |
| DPBS (1x) | Sigma-Aldrich | Cat# D8357 |
| DMEM (1x) | Sigma-Aldrich | Cat# D6546 |
| Immobilon® Western HRP Substrate | Merck Millipore | Cat# WBKLS0500 |
| Color Prestained Protein Standard, Broad Range (10-250 kDa) | New England BioLabs | Cat# P7719S |
| Rotiphorese.Gel 30 (37, 5:1) | Carl Roth | Cat# 3029.1 |
| TrypLE Express | Thermo Fisher Scientific | Cat# 12604013 |
| Trypsin | Promega | Cat# V5113 |
| Ammonium bicarbonate (ABC) | Fluka | Cat# 09830-100g |
| Streptavidin magnetic beads | New England BioLabs | Cat# S1420S |
| Carboxylate-coated magnetic beads (hydrophobic) | Cytiva | Cat# 65152105050250 |
| Carboxylate-coated magnetic beads (hydrophilic) | Cytiva | Cat# 45152105050250 |
| Monensin | ABCR | Cat# AB349208 |
| Bafilomycin | ABCR | Cat# AB355587 |
| Rapamycin | ABCR | Cat# AB352180 |
| TTFA | ABCR | Cat# AB104115 |
| 2-Deoxy-D-glucose | ABCR | Cat# AB173119 |
| 5/6-TAMRA-azide-biotin | Jena Bioscience | Cat# CLK-1048-5 |
| Fetal bovine serum | Thermo Fisher Scientific | Cat# A3840001 |
| Ethanol (EtOH) | Merck | Cat# 34852 |
| Chloracetamide (CAA) | Sigma-Aldrich | Cat# C0267-100g |
| pro-N6pA probe | / | (Kielkowski *et al*, 2020b) |
| GlcNAz probe | / | (Li *et al*, 2016) |
| **Software** |  |  |
| MaxQuant | Cox et al., 2014 | https://www.maxquant.org/download_asset/maxquant/latest |
| Perseus | Tyanova et al., 2016 | https://maxquant.net/download_asset/perseus/latest |
| Origin | N/A | https://www.originlab.com/ |
| **Other** |  |  |
| Pierce® BCA Protein Assay Kit | Thermo Fisher Scientific | Cat# 23225 |
| Orbitrap Eclipse Tribrid Mass Spectrometer | Thermo Fisher Scientific | N/A |
| FAIMS Pro Duo Interface | Thermo Fisher Scientific | N/A |
| PicoTip™ Emitter, Silica Tip™ | New Objectives | FS360-75-8-N-20-C15 |
| ReproSil-Pur 120 C18-AQ | Dr. Maisch GmbH | Cat# R119.aq.0001 |
| PEPMAP100 C18 5UM 0.3X5MM | Thermo Fisher Scientific | Cat# 160454 |

### Methods and Protocols

#### Culturing of HeLa and SH-SY5Y cells

HeLa (RRID: CVCL_0030) and SH-SY5Y (RRID: CVCL_0019) cells were cultured in Dulbeccos Modified Eagles Medium - high glucose (DMEM) supplemented with 10% fetal calf serum (FCS) and 2 mM L-alanyl-glutamine at 37°C and 5% CO_2_ atmosphere.

#### Probe and inhibitor treatment

In each 10 cm dish 2.5 million HeLa or SH-SY5Y cells were seeded in 10 mL media. Cells were either treated with 10 μL probe (100 mM stock pro-N6pA or 200 mM stock GlcNAz) or with 10 μL DMSO as a control. In case the cells were additionally treated with an inhibitor, the final concentrations were as follows: 100 nM rapamycin, 100 nM bafilomycin, 2 μM monensin, 100 μM TTFA and 4 μM 2-Deoxy-D-glucose. For the monensin concentration dependency experiments the final monensin concentrations were 2 nM, 20 nM, 200 nM, 1 μM and 2 μM. After addition of probe and inhibitor, the cells were incubated 16 h at 37 °C before harvesting. For this, the cells were washed twice with 2 mL DPBS, scrapped into 1 mL DPBS and pelleted at 260x g, 4 °C.

#### Cell lysis

Cells were lysed with 500 μL lysis buffer (20 mM Hepes, pH 7.5, 1% (v/v) NP40, 0.2% (w/v) SDS) and sonicated for 10 s at 20% intensity with a rod sonicator. Lysate was clarified with 12,000x g at 4 °C for 10 min and the protein concentration was determined by BCA. For optimization of different lysis buffers, following compositions were used: Lysis buffer 1 (8 M urea in 100 mM Tris/HCl pH 8.5) Lysis buffer 2 (1% NP-40 in PBS), Lysis buffer 3 (1% NP-40, 0.2% SDS in PBS), Lysis buffer 4 (0.5% Triton in PBS), Lysis buffer 5 (0.5% Triton, 0.2% SDS in PBS), Lysis buffer 6 (1% NP-40 in 20 mM Hepes pH 7.5), Lysis buffer 7 (0.5% Triton, 0.2% SDS in 20 mM Hepes pH 7.5), Lysis buffer 8 (0.5% Triton in 20 mM Hepes pH 7.5).

#### Measurement of protein concentrations

In order to measure the protein concentrations of the lysates, bicinchoninic acid assay was performed. First, bovine serum albumin (BSA) standards with concentrations of 12.5, 25, 50, 100, 200 and 400 μg/mL were prepared and samples as well as controls were diluted 40 times to a total volume of 200 μL. To measure standards, samples and controls in triplicates, 50 μL of each was added to three wells of a transparent 96-well plate with flat bottom. Afterwards, 100 μL working reagent (2 μL R2 and 98 μL R1) was added to each well by a multistepper and the plate was incubated 15 min at 60°C. Then, the absorbance at 620 nm was measured by Tecan and the protein concentrations were calculated.

#### SP2E workflow large scale

400 μg protein of probe treated and control lysates were diluted with lysis buffer (20 mM Hepes, pH 7.5, 1% (v/v) NP40, 0.2% (w/v) SDS) to 200 μL reaction volume. To each replicate, 2 μL biotin-PEG-N_3_ (10 mM in DMSO), 2 μL TCEP (100 mM in water) and 0.25 μL TBTA (83.5 mM in DMSO) were added. Samples were gently vortexed, click reaction was initiated by the addition of 4 μL CuSO_4_ (50 mM in water) and incubated for 1.5 h (rt, 600 rpm). Subsequently, 200 μL of 8 M urea was added to each replicate. 100 μL of mixed hydrophobic and hydrophilic carboxylate-coated magnetic beads (1:1) were washed thrice with 500 μL water. Click reaction mixture was directly transferred onto the equilibrated carboxylate-coated magnetic beads, resuspended and 600 μL ethanol was added. After resuspending the beads via vortexing, suspension was incubated for 5 min at rt and 950 rpm. The beads were washed thrice with 500 μL of 80% ethanol in water and the proteins were separately eluted by the addition of 0.5 mL of 0.2% SDS in PBS. For this, the beads were resuspended, incubated for 5 min at 950 rpm, rt and the supernatant was directly transferred onto 50 μL equilibrated streptavidin-coated magnetic beads (3 times pre-washed with 500 μL 0.2% SDS in PBS). The procedure was repeated once, the supernatants were combined and incubated for 1 h, rt and 950 rpm for biotin/streptavidin binding. The streptavidin-coated magnetic bead mixture was washed thrice with 500 μL 0.1% NP40 in PBS, twice with 500 μL 6 M urea and twice with 500 μL water. Washed bead mixtures were resuspended in 80 μL 125 mM ABC buffer and proteins were reduced and alkylated by the addition of 10 μL 100 mM TCEP and 10 μL 400 mM chloracetamide and 5 min incubation at 95 °C. Proteins were digested overnight at 37 °C with 1.5 μL sequencing grade trypsin (0.5 mg/mL). The following day, the beads were washed thrice with 100 μL 100 mM ABC buffer and the supernatants were combined and acidified with 2 μL formic acid. Peptides were desalted using 50 mg SepPak C18 cartridges on a vacuum manifold. The columns were equilibrated with 1 mL acetonitrile, 1 mL elution buffer (80% acetonitrile with 0.5% formic acid in water) and 3 mL wash buffer (0.5% formic acid in water). Subsequently, the samples were loaded on the cartridges and washed with 3 mL wash buffer. The peptides were eluted two times with 250 μL elution buffer and vacuum dried with a SpeedVac at 35 °C. Finally, dried peptides were reconstituted in 30 μL 1% formic acid in water by vortexing and sonication (15 min) and transferred to a MS vial.

During the optimization process, we performed this workflow with the following changes. The first attempts were performed without a separate elution step of the proteins from the carboxylated beads. In these attempts one was performed without the addition of 8 M urea after the click reaction. Furthermore, once the reduction and alkylation was performed prior to the click reaction. In addition, TEAB was used instead of ABC buffer for the digest. Another attempt was to reduce the starting protein amount from 400 μg to 100 μg.

#### SP2E workflow small scale

100 μg protein of probe treated and control lysates were diluted with lysis buffer (20 mM Hepes, pH 7.5, 1% (v/v) NP40, 0.2% (w/v) SDS) to 19 μL reaction volume in a 96-well plate. To each replicate, 0.2 μL biotin-PEG-N_3_ (10 mM in DMSO), 0.2 μL TCEP (100 mM in water) and 0.125 μL TBTA (16.7 mM in DMSO) were added. Samples were gently vortexed, click reaction was initiated by the addition of 0.4 μL CuSO_4_ (50 mM in water) and incubated for 1.5 h (rt, 600 rpm). Subsequently, 60 μL of 8 M urea was added to each replicate. 100 μL of mixed hydrophobic and hydrophilic carboxylate-coated magnetic beads (1:1) were washed thrice with 100 μL water. Click reaction mixture was directly transferred onto the equilibrated carboxylate-coated magnetic beads, resuspended and 100 μL of absolute ethanol was added. After resuspending the beads via vortexing, suspension was incubated for 5 min at rt and 950 rpm. The beads were washed thrice with 150 μL of 80% ethanol in water and once with 150 μL acetonitrile (LC-MS). Proteins were separately eluted by the addition of 60 μL of 0.2% SDS in PBS. For this, the beads were resuspended, incubated for 5 min at 40 °C and 950 rpm and the supernatant was directly transferred onto 50 μL equilibrated streptavidin-coated magnetic beads (3 times pre-washed with 100 μL 0.2% SDS in PBS). The procedure was repeated twice, the supernatants were combined and incubated for 1 h at rt and 800 rpm for biotin/streptavidin binding. The streptavidin-coated magnetic bead mixture was washed thrice with 150 μL 0.1% NP40 in PBS, twice with 150 μL 6 M urea and twice with 150 μL water. For each washing step, the beads were incubated 1 min at rt and 800 rpm. Washed bead mixtures were resuspended in 50 μL 50 mM TEAB and proteins were digested overnight at 37 °C by addition of 1.5 μL sequencing grade trypsin (0.5 mg/mL). The following day, the beads were washed twice with 20 μL 50 mM TEAB buffer and twice with 20 μL 0.5% FA and the wash fractions were collected and combined. For each washing step, the beads were incubated 5 min at 40 °C and 600 rpm. The combined washed fractions were acidified by addition of 0.9 μL formic acid (FA) and transferred to an MS-vial.

#### SPAAC protocol

400 μg protein of probe treated and control lysates were diluted with lysis buffer (20 mM Hepes, pH 7.5, 1% (v/v) NP40, 0.2% (w/v) SDS) to 200 μL reaction volume. To each replicate, 3 μL of 1M IAA in water was added and incubated for 30 min at 750 rpm, 25 °C. Next, 2 μL of 2 mM DBCO-PEG-N3 reagent was added to initiate SPAAC reaction. The reaction mixtures were incubated at 25 °C, 750 rpm for 30 min. The samples were proceeded further in the same way as for CuAAC.

#### MS-measurement

MS measurements were performed on an Orbitrap Eclipse Tribrid Mass Spectrometer (Thermo Fisher Scientific) coupled to an UltiMate 3000 Nano-HPLC (Thermo Fisher Scientific) via an EASY-Spray source (Thermo Fisher Scientific) and FAIMS interface (Thermo Fisher Scientific). First, peptides were loaded on an Acclaim PepMap 100 μ-precolumn cartridge (5 μm, 100 Å, 300 μm ID x 5 mm, Thermo Fisher Scientific). Then, peptides were separated at 40 °C on a PicoTip emitter (noncoated, 15 cm, 75 μm ID, 8 μm tip, New Objective) that was in house packed with Reprosil-Pur 120 C18-AQ material (1.9 μm, 150 Å, Dr. A. Maisch GmbH). The long gradient was run from 4-35.2% acetonitrile supplemented with 0.1% formic acid during a 150 min method (0-5 min 4%, 5-6 min to 7%, 7-105 min to 24.8%, 105-126 min to 35.2%, 126-140 min 80%, 140-150 min 4%) at a flow rate of 300 nL/min. The short gradient was run from 4-35.2% acetonitrile supplemented with 0.1% formic acid during a 60 min method (0-5 min 4%, 5-6 min to 7%, 7-36 min to 24.8%, 37-41 min to 35.2%, 42-46 min 80%, 47-60 min 4%) at a flow rate of 300 nL/min. FAIMS was performed with two alternating CVs including −50 V and −70 V. For measurements of chemical-proteomics samples, the Orbitrap Eclipse Tribrid Mass Spectrometer was operated in dd-MS^2^ mode with following settings: Polarity: positive; MS^1^ resolution: 240k; MS^1^ AGC target: standard; MS^1^ maximum injection time: 50 ms; MS^1^ scan range: m/z 375-1500; MS^2^ ion trap scan rate: rapid; MS^2^ AGC target: standard; MS^2^ maximum injection time: 35 ms; MS^2^ cycle time: 1.7 s; MS^2^ isolation window: m/z 1.2; HCD stepped normalised collision energy: 30%; intensity threshold: 1.0e4 counts; included charge states: 2-6; dynamic exclusion: 60 s.

#### Quantification and statistical analysis

MS raw files were analysed using MaxQuant software 2.0.1.0 with the Andromeda search engine. Searches were performed against the Uniprot database for Homo sapiens (taxon identifier: 9606, March 2020). At least two unique peptides were required for protein identification. False discovery rate determination was carried out using a decoy database and thresholds were set to 1% FDR both at peptide-spectrum match and at protein levels. LFQ quantification was used as described for each sample. For calculation of the large scale samples, carbamidomethylation was set as a fixed modification and methionine oxidation as well as N-terminal acetylation as a variable modification. In contrast, the small scale samples of the 96-well plate were calculated without carbamidomethylation as a fixed modification.

Statistical analysis of the MaxQuant result table proteinGroups.txt was done with Perseus 1.6.10.43. First, LFQ intensities were log_2_-transformed. Afterwards, potential contaminants as well as reverse peptides were removed. Then, the rows were divided into two groups - DMSO (control) and probe treated sample (sample). Subsequently, the groups were filtered for at least three valid values out of four rows in at least one group and the missing values were replaced from normal distribution. The -log_10_(*p*-values) were obtained by a two-sided one sample Student’s t-test over replicates with the initial significance level of p = 0.05 adjustment by the multiple testing correction method of Benjamini and Hochberg (FDR = 0.05) using the volcano plot function.

#### Western blot analysis

For each Western blot analysis, 20 μg cell lysate was used. In order to denature proteins, 4 μL 5× Laemmli buffer (10% (w/v) SDS, 50% (v/v) glycerol, 25% (v/v) β-mercaptoethanol, 0.5% (w/v) bromphenol blue, 315 mM Tris/HCl, pH 6.8) was added to 16 μL lysate solution and the samples were boiled 5 min at 95 °C. Afterwards, 20 μL of each sample was loaded onto a 7.5, 10 or 12.5% SDS gel and proteins were separated according to their size by SDS-PAGE. Then, the separated proteins were transferred onto a membrane using a blotting sandwich moistened by blotting buffer (48 mM Tris, 39 mM glycine, 0.0375% (m/v) SDS, 20% (v/v) methanol), which was composed of one extra thick blot paper, the PVDF transfer membrane, the SDS-PAGE gel and again one extra thick blot paper. Before the protein transfer was carried out 45 min at 25 V using a Semi Dry Blotter (Bio-Rad), the transfer membrane was pre-incubated 5 min in methanol. In order to block non-specific binding sites, the membrane was incubated 60 min in blocking solution (0.5 g milk powder in 10 mL PBST (PBS +0.5% Tween)). Subsequently, 10 μL primary antibody with specificity for the protein of interest was added and the mixture was incubated 1 h at room temperature. The membrane was washed 3 times for 10 min with PBST before 1 μL of the secondary HRP antibody in 10 mL blocking solution was added. After 1 h of incubation at room temperature, the membrane was washed again 3 times for 10 min with PBST. Then, 400 μL ECL Substrate and 400 μL peroxide solution were mixed and added to the membrane to stain the Western blot. Finally, images of the Western blot were taken by developing machine Amersham Imager 680 (GE Healthcare).

#### Trifunctional linker enrichment

400 μg protein of probe treated and control lysates were diluted with lysis buffer (20 mM Hepes, pH 7.5, 1% (v/v) NP40, 0.2% (w/v) SDS) to 200 μL reaction volume. To each replicate, 2 μL 5/6-TAMRA-azide-biotin (10 mM in DMSO), 2 μL TCEP (100 mM in water) and 0.25 μL TBTA (83.5 mM in DMSO) were added. Samples were gently vortexed, click reaction was initiated by the addition of 4 μL CuSO_4_ (50 mM in water) and incubated for 1.5 h (rt, 600 rpm). The clean-up of the click reaction mixture on the carboxylate magnetic beads as well as the subsequent enrichment on the streptavidin magnetic beads was performed as described in the above section of the SP2E large scale protocol. The enriched proteins were eluted from the beads by addition of 50 μL 1x Laemmli buffer and 5 min incubation at 95 °C. Finally, 20 μL of the eluate were loaded on a SDS-gel and the western blot was performed as described in the above section.

## Data Availability

Proteomics data are freely available at ProteomeXchange Consortium via PRIDE partner repository with the dataset identifier PXD030960.

## Acknowledgements

This research project was supported by Liebig fellowship from VCI to PK and TB, LMUexcellence Junior Fund to PK and SFB1309 by DFG.

## Author contribution

PK designed the method; AW, AT and DB performed the experiments and optimization of the method; TB carried out the AMPylation study; A H.-R. provided the GlcNAz probe and experience in *O*-GlcNAcylation. TB and PK overviewed the study and analyzed the data; TB and PK wrote the manuscript with input from all authors.

