## Supplementary Information for "Transforming chemical proteomics enrichment into high-throughput method using SP2E workflow"

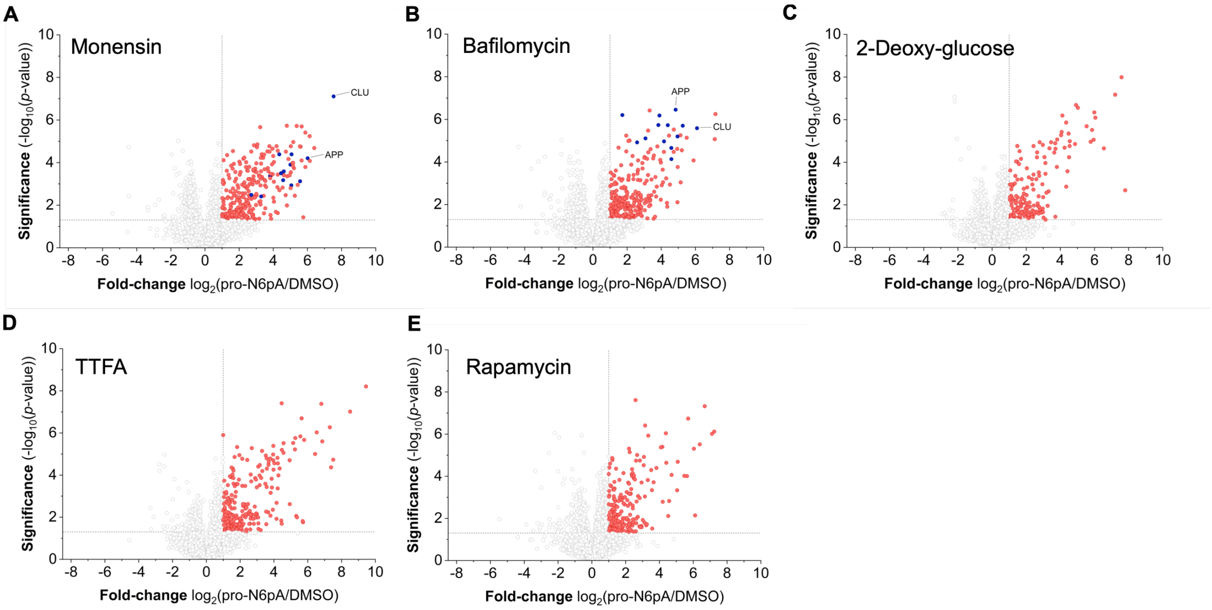


**Figure S1. A-E** Volcano plots showing the enrichment of pro-N6pA modified proteins in cell cultures treated with different inhibitors. $n$ = 4, cut-off lines at $p$-value >0.05 and 2-fold enrichment. Red dots are significantly enriched proteins. Blue dots represent proteins with a similar profile plot to APP, which were found significantly enriched only after monensin and bafilomycin treatment. This group includes following proteins: GPR56, FAT1, LAMA4, TGOLN2, RNF149, CRIM1, ITM2B, L1CAM, TMEM59, MCAM, LRP1 and CLU.

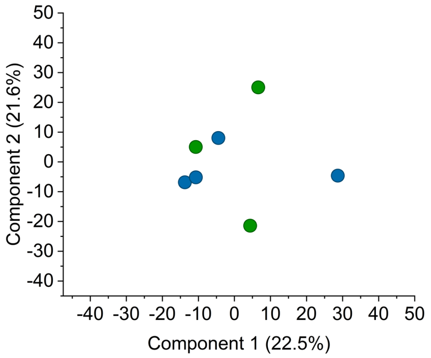


**Figure S2.** PCA of LFQ intensities from momensin treated cells with addition of DMSO (blue dots) or pro-N6pA probe (green dots) after subtraction of significantly enriched proteins (Fig S1A). For comparison see Fig 3C.

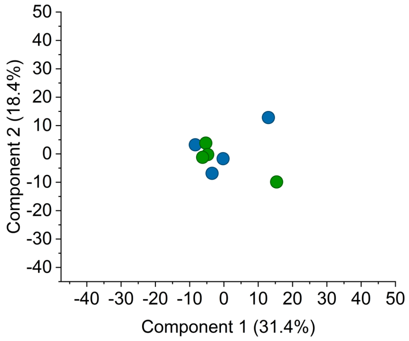


**Figure S3.** PCA of LFQ intensities from GlcNAz probe treated cells (green dots) or DMSO (blue dots) after subtraction of significantly enriched proteins (Fig 5B). For comparison see Fig 5C.

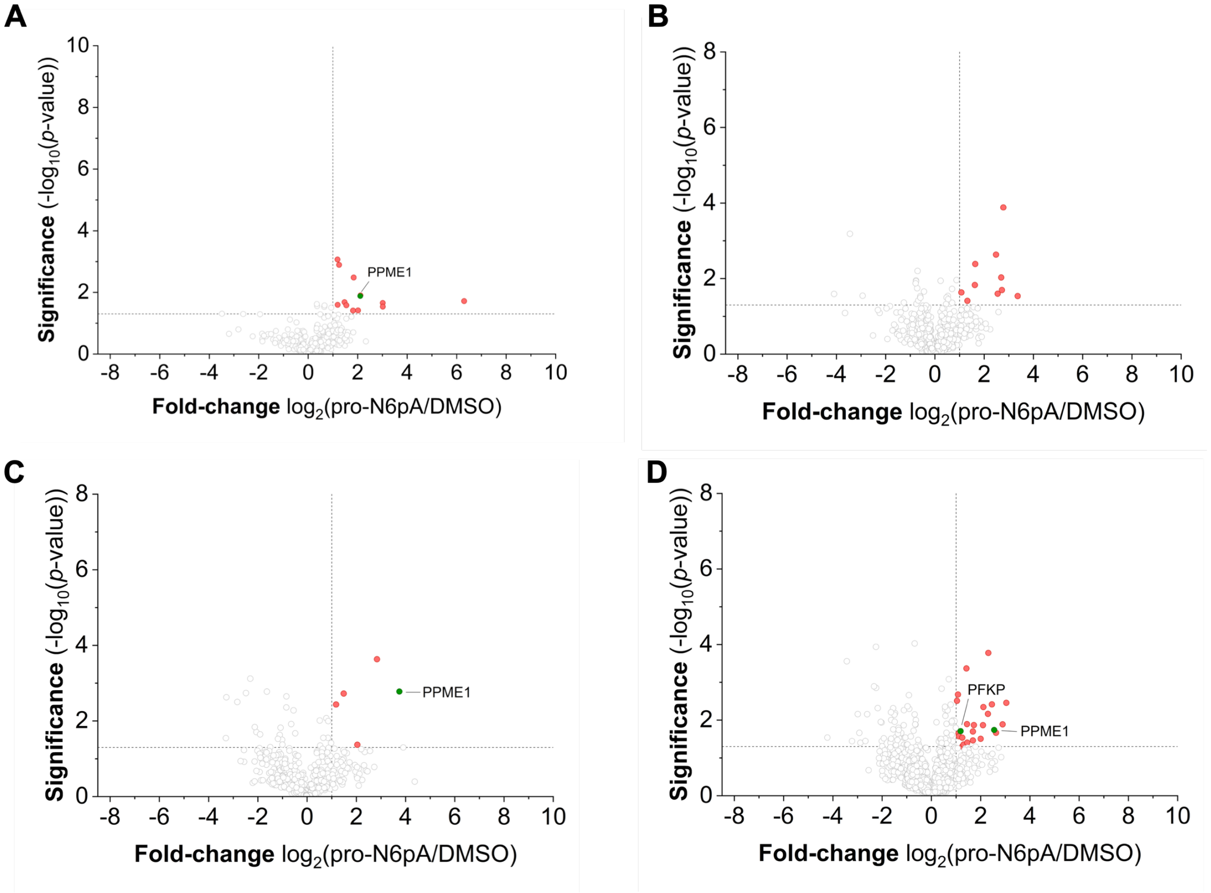


**Figure S4.** Volcano plot showing the optimization of the 96-well plate format SP2E workflow starting with 100 µg total protein. All samples have been analysed using the 150 min LC-MS/MS gradient. For comparison see Fig 6B. Following is description of the alternations to the protocol described in Methods and Materials. **A)** The clean-up of the proteins without ACN wash step and proteins were digested in the ABC buffer. **B)** The clean-up of the proteins without ACN wash step. Protein digest was carried out in ABC buffer. Additional elution with 1% FA after proteins' digest was omitted to decrease the dilution of the peptides. **C)** The amounts of carboxylate magnetic and streptavidin magnetic beads was halved. The clean-up of the proteins without ACN wash step and proteins were digested in the ABC buffer. **D)** Additional elution with 1% FA after proteins' digest was omitted to decrease the dilution of the peptides.

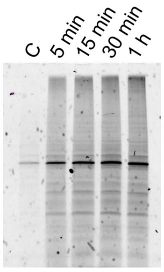

**Figure S5.** Biotin-streptavidin complex formation – incubation time optimization.
